# Probing the hierarchical dynamics of DNA-sperm nuclear transition protein complex through fuzzy interaction and mesoscale condensation

**DOI:** 10.1101/2023.02.14.528413

**Authors:** Shangqiang Xie, Congran Yue, Sheng Ye, Zhenlu Li

**Affiliations:** School of Life Science, Tianjin University, 92 Weijin Road, Tianjin 300072, China; Frontiers Science Center for Synthetic Biology (Ministry of Education), Tianjin Key Laboratory of Function and Application of Biological Macromolecular Structures, Tianjin University, 92 Weijin Road, Tianjin 300072, China

## Abstract

Nuclear transition protein TNP1 is a crucial player mediating histone-protamine exchange in condensing spermatids. A unique combination of intrinsic disorder and multivalent properties turns TNP1 into an ideal agent for orchestrating the formation of versatile TNP-DNA assembly and endows the protein with potent value for vaccine design. Despite its significance, the physicochemical property and the molecular mechanism taken by TNP1 for histone replacement and DNA condensation are still poorly understood. In this study, for the first time, we expressed and purified in vitro human TNP1. We investigated the hierarchical dynamics of TNP1: DNA interaction by combing computational simulations, biochemical assay, fluorescence imaging, and atomic force microscopy. We analyzed fuzzy interactions between TNP1 and DNA at the atomistic level and assessed the influence of TNP1 association on the electrostatic and mechanical properties of DNA. Furthermore, the alteration of the physicochemical properties of the TNP1-DNA complex modulates its molecular assembly and phase separation. Our study sets the foundation for understanding TNP1-mediated histone replacement and sheds light on the encapsulation of genetic material by TNP1 for vaccine development.

## Introduction

Spermatogenesis is a process by which haploid spermatozoa develop into mature sperm cells with suitable functions to meet and interact with the egg (Brinster and Zimmermann, 1994). During the late stage of spermatogenesis, the cell nucleus flattens and condenses. One of the major changes in the nucleus is the disassembly of histone from the nucleosome, and subsequently, the genetic material DNA is compressed into a highly condensed form (Braun, 2001; Brewer et al., 2002; Kota and Feil, 2010). This process completely shutdowns the transcription of the gene, and thus the genetic material could be well protected during the passing on of genetic information (Aoki et al., 2006; Gou et al., 2020; Merges et al., 2022). The removal of histone and the condensation of DNA is facilitated by several basic disordered proteins, of which Spermatid nuclear transition protein (TNP1 and TNP2) and sperm protamine (P1 and P2) are currently best known (Balhorn, 2007; Meistrich et al., 2003). These proteins are highly positively charged and thus can compete with and replace histone for wrapping and encapsulating DNA.

Nuclear transition protein and sperm protamine are of great significance for the safe passing of genetic material. Additionally, due to their unique DNA-binding ability, these kinds of small basic proteins have great application prospects in the field of vaccine design. Protamine from salmon fishes, in particular, has been explored in academia and in the industry as a tool to encapsulate pharmaceutical nucleic acid substances and deliver them to cells (Mai et al., 2020; Ruseska et al., 2021; Scheel et al., 2005). Similar types of nucleic acids-condensing proteins such as PEG10 and phase-separated peptides derived from histidine (His)-rich beak proteins (HBPs) have recently been developed as promising tools for mRNA nanomedicine (Huang X et al., 2022; Segel et al., 2021; Sun et al., 2022).

Despite their significance in the preservation of genetic material and great potency in vaccine development, the detailed molecular mechanism taken by the nuclear transition protein and sperm protein for histone replacement and DNA condensation is still largely unknown. The multibody interactions between histone, DNA and the disordered basic proteins reflect higher-order dynamical effects that include fuzzy interactions between intrinsic disorder protein and DNA (Turner et al., 2018), biomolecular condensation and phase separation (Gou et al., 2020; Kang et al., 2022), as well as regular or irregular morphological changes (Gibson et al., 2019). These dynamical effects are spatiotemporally dependent and are regulated by multiple selves and environmental factors, such as the ratio of concentration and charge between molecules (DeRouchey et al., 2013; DeRouchey and Rau, 2011; Makita et al., 2011). The complicated spatiotemporal dynamics of basic proteins and nucleotide acids build a barrier for revealing the inherent mechanism (Heidarsson et al., 2022). And a comprehensive understanding of the process of molecular competing and assembly calls for dedicated characterization of molecular interaction at both microscopic and mesoscopic scales.

Of the four above-mentioned basic proteins that bind and condensate chromatin DNA, TNP1 and TNP2 are supposed to be the early molecules that bind to DNA and assist in the histone-protamine exchange (Gou et al., 2017; Lewis et al., 2003). Transition protein 1 (TNP1 or TP1) is an intrinsic disorder protein with abundant structural flexibility. A striking feature of TNP1 is the extensive use of arginine and lysine for DNA association. Of the 55 constituent amino acids of TNP1, there are 11 arginines, 10 lysines, and 2 aspartic acids, yielding a total of 19 net positive charges (Fig. 1a). This unique combination of intrinsic disorder and the multivalent properties turns TNP into ideal agents to orchestrate the formation of versatile TNP-DNA assembly in condensing spermatids, but it also challenges the in vitro expression and purification of TNP1. As far as we know, the purification of human source TNP1 with recombinant DNA techniques has been not reported in the literature, needless to say the full characterization of biochemical property. By fusing soluble proteins to TNP1 and making use of ion exchange columns and centrifugation, for the first time we successfully isolated human TNP1 (Fig. S1).

**Figure 1:**
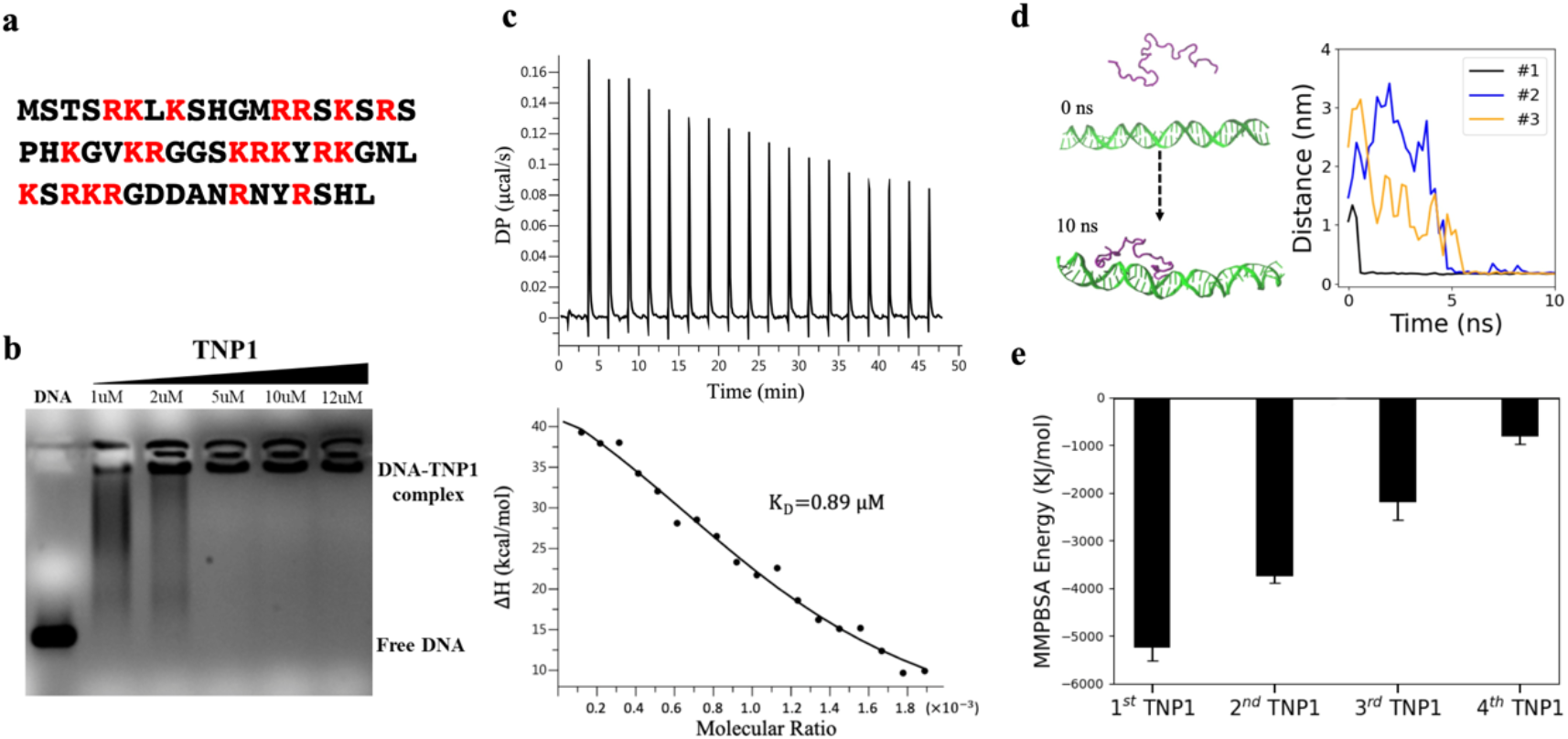
Associations of TNP1 with DNA. (a) TNP1 sequence. (b) Gel Electrophoresis of DNA in the presence of TNP1 at various concentrations. The DNA (177 bp) concentration is 450 uM/bp. (c) ITC binding studies between TNP1 and DNA. 0.2 uM TNP1 is titrated into a DNA solution at 2 uM/bp. (d) Time evolution of the minimal distance between TNP1 and DNA in MD simulations: The left images indicate the starting and a binding configuration (MD snapshot taken at 10 ns) of the TNP1 (purple) and DNA (green) complexes. (e) Estimations of MM-PBSA energy for sequentially binding of the first, second, third, and fourth TNP1 to a DNA molecule (40 bp).

With purified TNP1, we aimed to uncover the multiscale dynamic and assembly behaviors of TNP1 and DNA molecules at varying conditions. To achieve this goal, we employed a combination of biochemical assays, fluorescence imaging, single-molecule atomic force microscopy, and atomistic and mesoscopic molecular dynamics simulations. We examined the changes in the mechanical and electrical properties of DNA and analyzed the biomolecular condensation and crosslinking behavior. This study paves the way for a complete understanding of the assembly of genetic material by TNP under physiological conditions.

## Results and discussion

### Association of TNP1 with DNA

We firstly analyzed the gel electrophoresis of DNA in the presence of TNP1 at various concentrations (Fig. 1b). The ratio of the positive charge owned by proteins (19 |e|/TNP1) and the negative charges possessed by DNA (−2|e|/bp) --- R_+/−_, is a crucial determinant for the protein-DNA association. For TNP1, the gel electrophoresis assay shows that a small amount of TNP1 at 1 *μ*M (R_+/−_=0.02) can dramatically alter the mobility of DNA, leading to dispersed DNA migration bands. When TNP1 was added to 5 *μ*M, i.e., R_+/−_=0.1, it fully blocks the migration of DNA. The gel electrophoresis revealed the binding propensity between TNP1 and DNA. To further quantify the TNP1-DNA binding affinity and thermodynamic parameters, Isothermal Titration Calorimetry (ITC) was performed by titrating TNP1 into DNA solution. Fig. 1c shows that the TNP1-DNA association is endothermic with enthalpy ΔH of 56.1 kcal/mol. The enthalpy loss is compensated by a larger entropy contribution (-TΔS) of −64.3 kcal/mol, combing into a Gibbs free energy of −8.26 kcal/mol. The corresponding binding affinity K_d_ was measured at 0.89*μ*M. The entropy-enthalpy compensation is in trend consistent with the binding of DNA with proteins of the other types (Gupta et al., 2019; Jen-Jacobson et al., 2000).

Conventional molecular dynamics simulations were applied to further study the binding process of TNP1 toward DNA. In three independent simulations (Table S1), a free TNP1 was observed to rapidly bind to a DNA molecule within 10 ns (Fig. 1d). We further simulated the sequential binding of the 2^nd^, 3^rd^, and 4^th^ TNP1 toward the same DNA molecule, in order to investigate the influence of TNP1/DNA ratio. The association energy between the DNA and sequentially bound TNP1 was estimated with the MM/PBSA (Molecular Mechanics**/**Poisson-Boltzmann Surface Area) method. The MM-PBSA estimation for this kind of highly charged system usually lacks accuracy at the quantitatively level. However, at least at the qualitative level, the analysis presents a rapid rising energy landscape as the sequential addition of the 1^st^, 2^nd^, 3^rd^, and 4^th^ TNP1 (Fig. 1e and Figure S2), indicating weaker and weaker DNA-binding propensity of TNP1 as its content increases.

### Fuzzy interactions between TNP1 and DNA

Interactions involving disordered proteins often exhibit versatile binding modes and form fuzzy complexes. From the above simulations, we have learned that TNP1 easily binds to DNA. However, a system of interest is often trapped in metastable states in the conventional simulations, and hence it is difficult to efficiently sample conformational ensemble. Therefore, we further performed metadynamics simulations with 20 independent sets of constructs to explore the potent configurations of the TNP1-DNA complex (see Methods). The results indicate that TNP1 exhibit multiple configurations with different radius of gyration (R_g_) varying from 1.5 nm to 3 nm. The representative configurations include a stretched configuration with a large R_g_, a contracted configuration with small a R_g_, as well as configurations in between (Fig. 2a). These configurations relate to distinct binding modes toward DNA (Fig. 2a and Fig. S3). For a stretched TNP1, it is linearly anchored to DNA, while for a contracted TNP1, it tends to encompass and wrap DNA in whole or at least in part.

**Figure 2:**
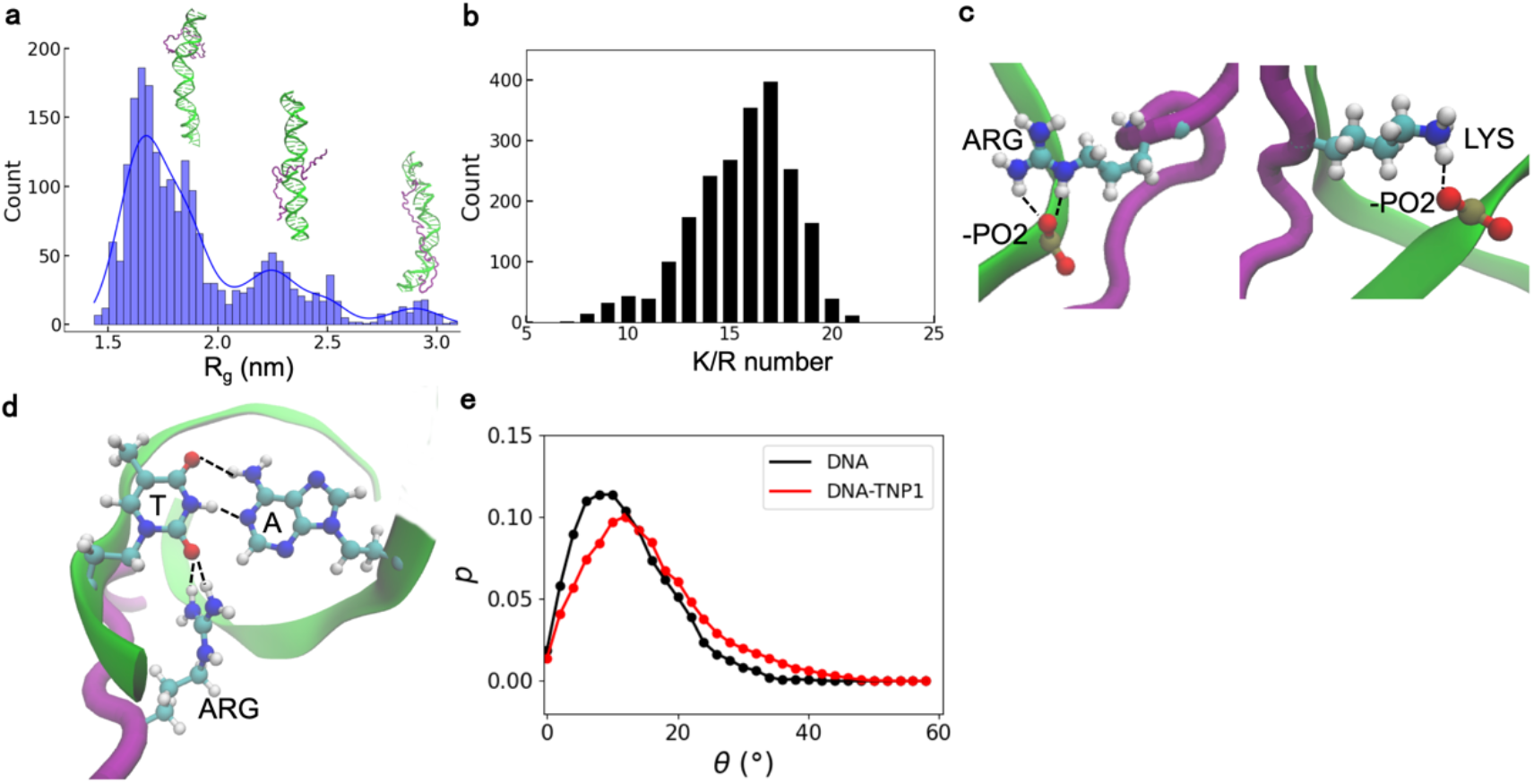
Fuzzy interactions between TNP1 and DNA. (a) Distribution of radius of gyration of TNP1: (b) Distribution of number of K/R bound to DNA. (c) Arginines and lysines use the guanidinium and amine group respectively to bind to the phosphate group (PO_2_-). (d) Groove insertion of arginine and establishment of interaction between guanidinium and the bare O3’ atom of thymine. (e) Computational analysis of the distribution of bending angle of DNA in the absence or presence of TNP1.

The median number of lysine and arginine (K/R) bound to DNA is counted as 17 (Fig. 2b). Thus, the majority of positively charged amino acids (totally 21 K/R) contribute to the DNA binding. In most cases, arginines and lysines bind to the phosphate group (PO_2-_) with the guanidinium and amine group respectively(Crothers et al., 1990; Rohs et al., 2009; Tesei et al., 2017). The guanidinium-phosphate group (PO_2-_) interactions are typically stronger than the amine-phosphate group interactions as the guanidinium exhibits multiple polar hydrogens enabling the coexistence of 2 hydrogen-bonds bridges between an arginine and a phosphate group (Rohs et al., 2009). In addition to the attraction between K/R and phosphate group, interestingly, we found that the guanidinium of arginine also interacts with oxygen at the 2’ position of thymine (Fig. 2d). For thymine, the NH at 3’ and O at 4’ position form two hydrogen bonds with adenine. However, the O2’ atom is unpaired and free. Thus the guanidinium of arginine could insert into the DNA groove and establish interaction with this oxygen.

On the side of DNA, in all the 20 sets of simulations, DNA bends at varying extents (Fig. S3). By calculating the angle between any 3 connected groups each consisting of 10 bps (see definition in Fig. S4), we found that the association of TNP1 increased the defined bending angle of DNA (Fig. 2e). In some area, especially at the location of TNP1 binding, the bending angle can even reach 30 to 40 degrees.

### Influence of TNP1 on electrostatic and mechanical properties of DNA

The association of TNP1 with DNA may alter the structural, electrostatic and mechanical properties of DNA. The zeta potential demonstrates an indication of the surface charge on a particulate species. In Fig. 3a, the zeta potential of DNA was measured against increasing TNP1 concentration. The result indicated pure DNA has a negative zeta potential about −21 mV. The addition of TNP1 of 1 or 5 *μ*M (R_+/−_=0.05-0.25) reduces the zeta potential to −15 mV. Further addition of TNP1 (10-15 *μ*M, R_+/−_=0.5-0.75) brings the zeta potential to about −5 mV level. In addition to the neutralization effect, the TNP1: DNA attraction may also potentially cause changes in the DNA secondary structure(Gupta et al., 2019). To ascertain potent structural changes, CD spectra of DNA at pH 7.4 were recorded and found to be very different in the presence and absence of TNP1. The addition of TNP1 affects the intensity of the CD spectrum. As Figure 3b shows, the lower peak at about 230 nm is sharp in the presence of TNP1. The positive peak near 275 nm is increased and shifted with the addition of TNP1 of 1 or 2.5 *μ*M. However, the peak intensity is decreased when the TNP1 concentration increases to 10 *μ*M. These peak changes potentially relate to the observed deformation of DNA in simulation under the influence of TNP1.

**Figure 3:**
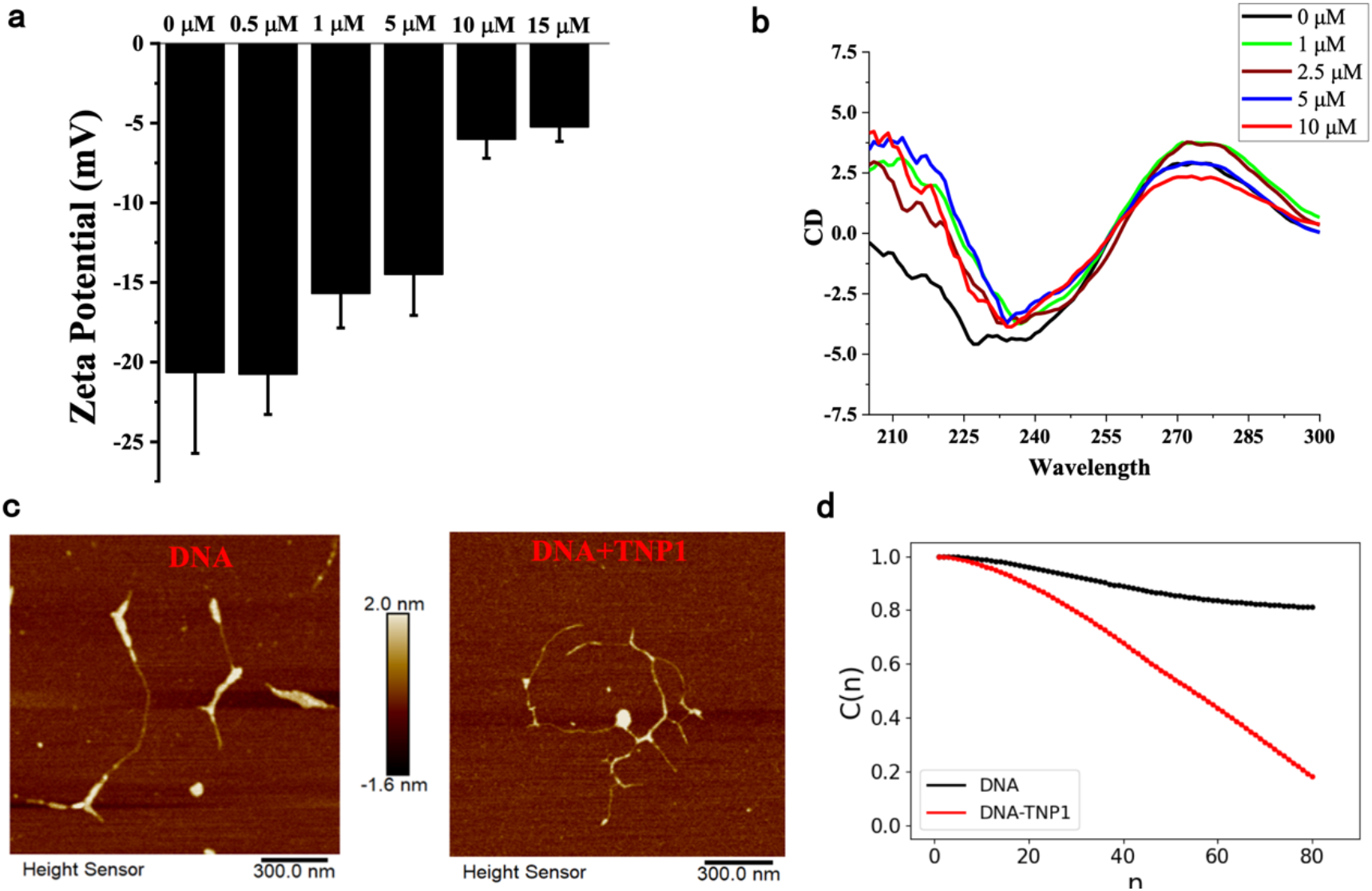
Influence of TNP1 on electrostatic and mechanical properties of DNA. (a) Zeta potential of DNA (75 μM/bp) in the presence of TNP1 at varying concentrations. (b) CD spectrum of DNA (450 μM) under the influence of TNP1 at different concentrations. (c) AFM images of pure DNA (left, 1.5 μM/bp) and DNA-TNP1 complex (right, DNA of 0.3 μM/bp and TNP1 of 0.1 μM). (d) Autocorrelation function C(n) as a function of distance between two DNA segments (d=n×d_l_, d_l_ = 8.1 nm).

To further assess the changes in mechanical properties of DNA due to TNP1 association, we used atomic force microscopy (AFM) to illustrate and compare the morphology of a long DNA molecule (3043 bp) with and without TNP1 (Fig. 3c). The AFM imaging clearly shows a difference between pure DNA and the DNA-TNP1 mixture. A plain DNA is predominantly linear in shape. In comparison, the DNA-TNP1 mixture exhibit a very obvious looping and inter-strand crosslinking phenomenon (Fig. 3c and Fig. S5). By extracting the DNA trajectories from AFM images (Fig. S6), we further calculated the autocorrelation function C(n) as a function of distance n (See Methods). The analysis shows that the TNP1 association largely attenuates the correlations within a DNA molecule, reducing the stiffness of a DNA chain.

The above analysis indicates that TNP1 reduces the zeta potential of DNA and increases the flexibility of DNA molecules. As the zeta potential reduces and mechanical strength changes, the TNP1 and DNA complex would present higher-order dynamical effects such as condensation or phase separation.

### TNP1-DNA assembly at different ratios of TNP1 and DNA

To understand the molecular details of the multibody behavior of TNP1 and DNA mixtures, we simulated the TNP1-DNA assembly with a Weeks-Chandler-Anderson (WCA) potential-based coarse-grained model. This method has been used to study the assembly mechanism of salmon protamine and short DNA(Toma et al., 2009; Ukogu et al., 2020). Specifically, in five different simulations, 30 DNAs (40 bp in length) were mixed with 30, 60, 90, 120 and 180 TNP1 molecules, respectively. The corresponding R_+/−_ varies from 0.3 to 1.4. The results show that TNP1 could promote the formation of DNA bundles (Fig. 4a). Since TNP1 is unstructured and flexible, TNP1 can bind to one DNA with partial amino acids, and simultaneously bind to another DNA with the remaining amino acids. Therefore, TNP1 can bridge two DNAs. However, the bridging effect is counteracted by the electrostatic repulsion between DNA molecules.

**Figure 4:**
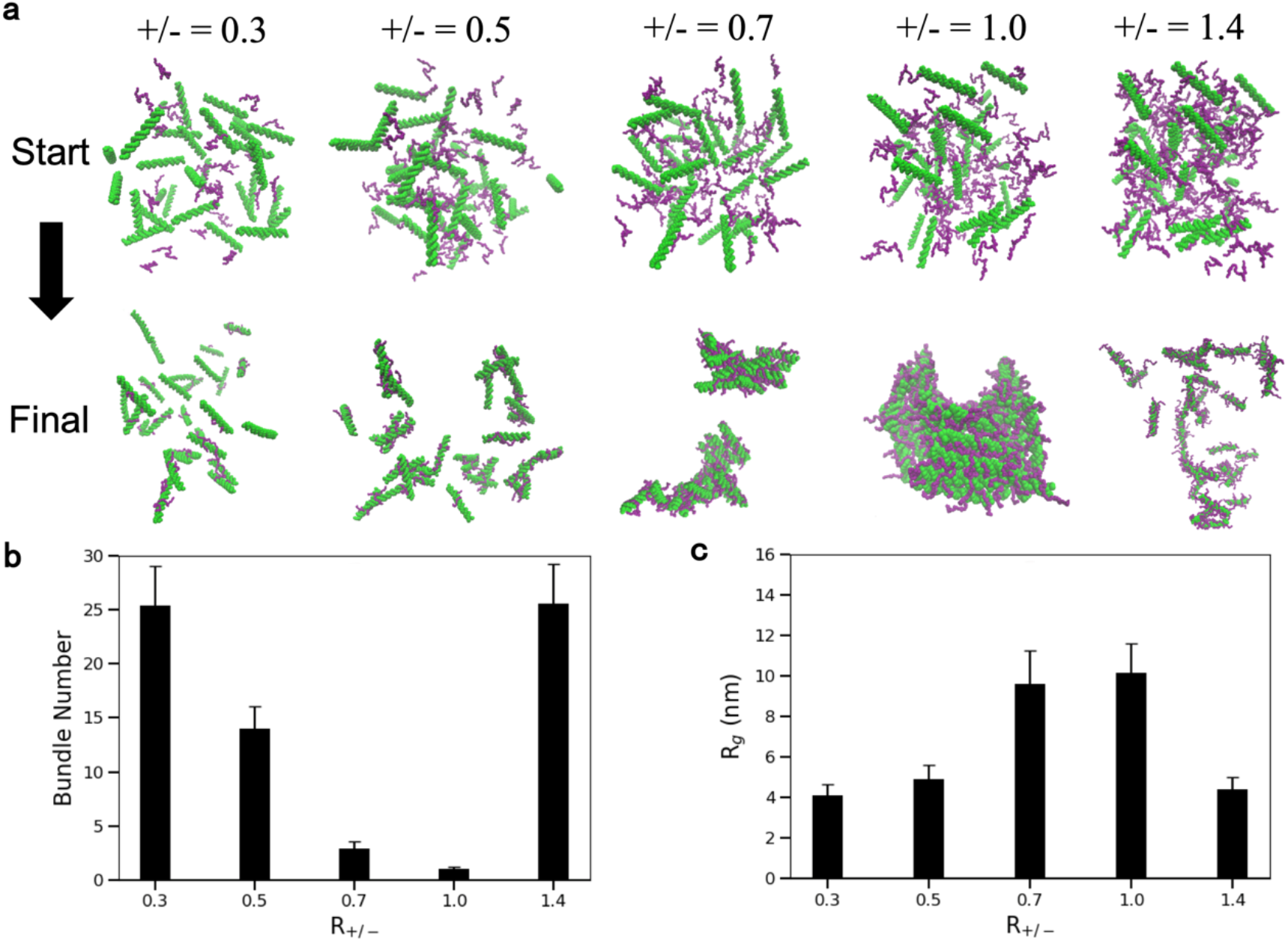
TNP1-DNA assembly at different R+/− ratios. (a) Simulations of the mixture of 30 DNA with 30, 60, 90, 120, and 180 TNP1. Thu upper images show the starting simulation configuration. The lower images show the final snapshots of the TNP1-DNA mixture. (b) Number of independent bundles of the TNP1-DNA mixture at different R_+/−_. (c) Weighted mean of radius of gyrations (R_g_) of all bundles at different R_+/−_.

At low contents of TNP1 (R_+/−_=0.3), the majority of DNA is monodisperse as exhibited by a large number of independent bundles (Fig. 4b) and a small value of radius of gyration (*R*_g_, weighted mean of the size of all bundles) (Fig. 4c). As the increase of TNP1 (R_+/−_=0.5), the electrostatic shielding effect of TNP1 enhances and larger bundles of 2 to 6 DNA molecules grow up. At higher content of TNP1 (R_+/−_=0.7 or R_+/−_=1.0), all DNA molecules are clustered into one or two big aggregates (Fig. 4b) and the average R_g_ increases up to 10 nm (Fig. 4c). Interestingly, when excess TNP1 were added, the TNP1-DNA mixture returns to a disperse state with only small bundles. These kinds of reversibly assembly and disassembly phenomenon have been observed in protamine-mediated DNA assembly as well as RNA-peptide/protein condensates (Banerjee et al., 2017; Henninger et al., 2021; Mukherjee et al., 2021a).

### TNP1-DNA condensation at high ratio of TNP1 to DNA

To further understand TNP1-DNA complex condensation, we again used AFM to illustrate the morphology of DNA under the influence of TNP1 at high concentrations. It was already shown in Fig. 3c that the DNA-TNP1 mixture exhibit an obvious looping and inter-strand crosslinking phenomenon at medium concentrations of TNP1 (Fig. 3c and Fig. S5). As the contents of TNP1 increase, the DNA-TNP1 mixture forms a denser hinged network, with some of the DNA compressed into microspheres (Fig. 5a). Dynamic light scattering was applied to measure the average size of the DNA and DNA-TNP1 mixture. At medium TNP1/DNA ratio (R_+/−_=0.5), most TNP1-DNA was mono-dispersed compared with pure DNA (Fig. 5b). As TNP1/DNA ratio increases (R_+/−_=1.0 or R_+/−_=2.0), obviously a peak of particle size of several micrometer shows up, indicating the appearance of larger particles in the mixture (Fig. 5b).

**Figure 5:**
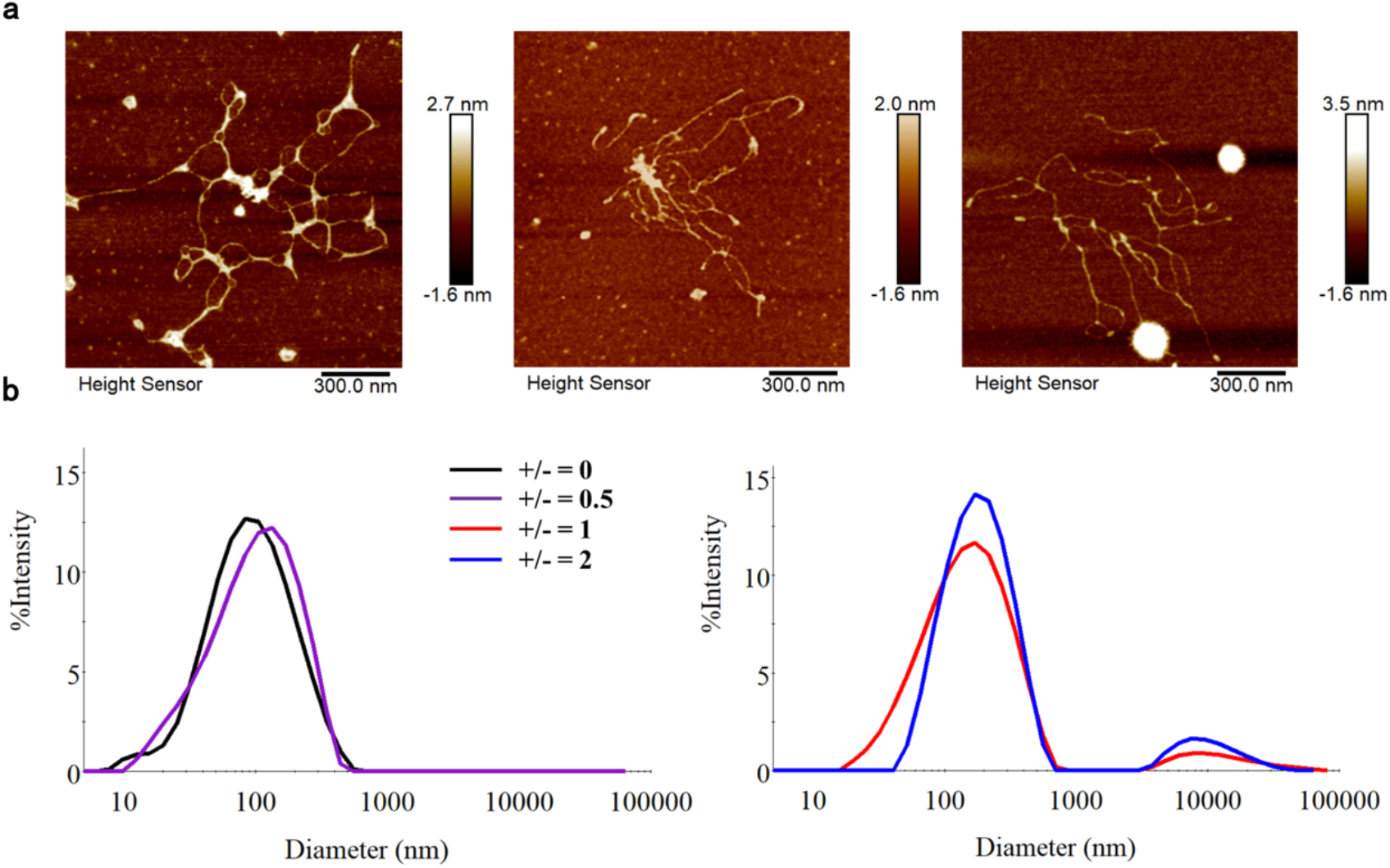
TNP1-DNA condensation. (a) AFM of TNP1-DNA mixture at high TNP1/DNA ratio (DNA of 0.3 μM/bp and TNP1 of 1.0 μM). Three representative fields of view. (b) Dynamic light scattering of DNA at varying concentration of TNP1 (DNA at 100 μM/bp).

### Formation of phases-separated droplets and potent influence factors

We hypothesized that the TNP1-DNA complex underwent higher-order dynamic change at various TNP and DNA ratios. We further expressed the eGFP-TNP1 fusion protein to observe the phase behavior of the eGFP-TNP1/DNA complex at various conditions. Firstly, we performed a control experiment by mixing 10 *μM* eGFP-TNP1 with positively charged arginine, negatively charged glutamine, as well as ATP molecules at various concentrations (10-100*μM*). Fig. 6a-b shows representative fluorescent images of eGFP-TNP1 solution and a mixed solution of eGFP-TNP1 and ATP molecules. In all these experiments, eGFP-TNP1 were evenly distributed in the solution and were not able to be gathered into phased-separated droplets.

**Figure 6:**
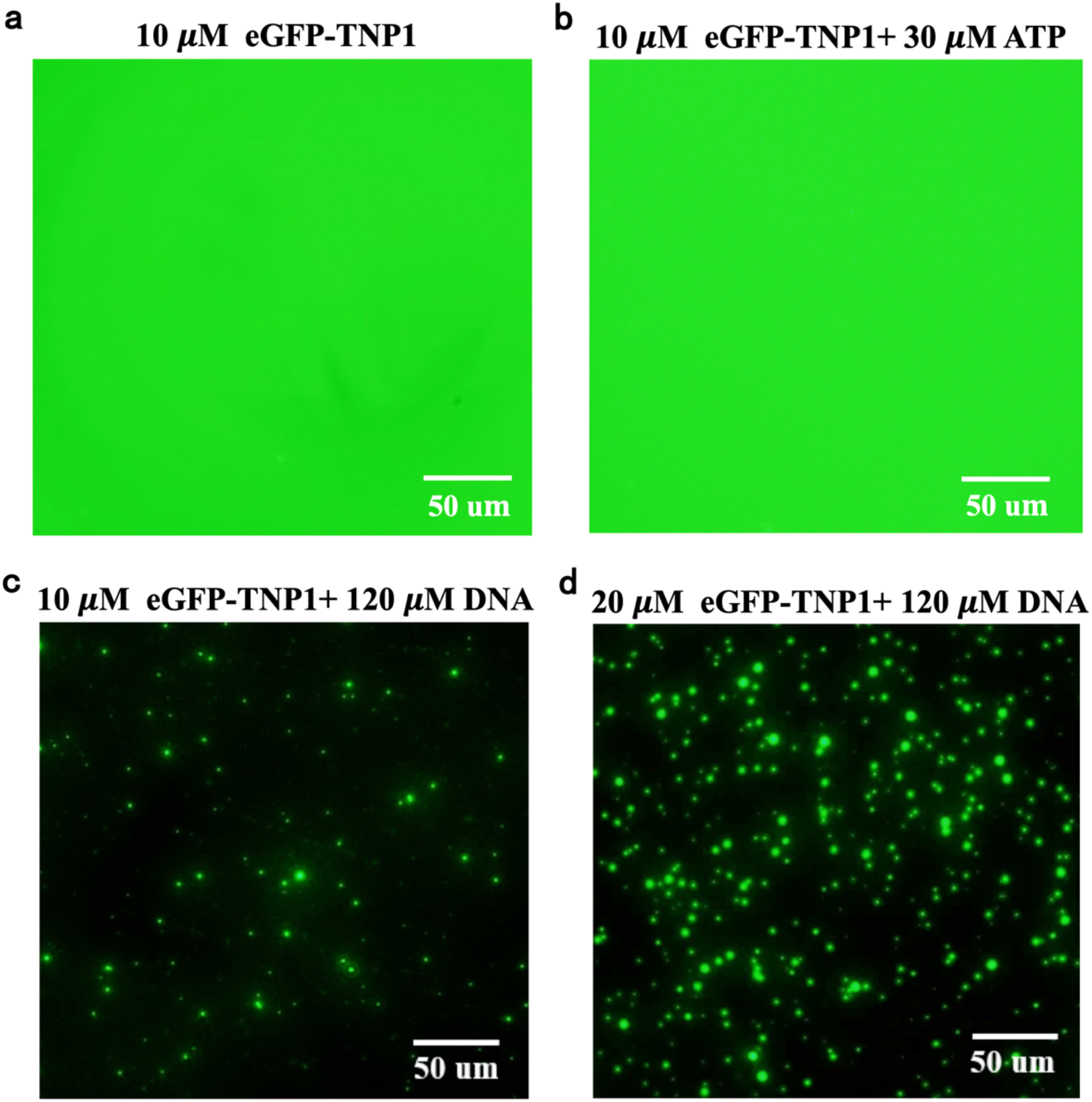
The formation of phases-separated droplets. Fluorescence imaging of (a) eGFP-TNP1 (10 μM) and (b) eGFP-TNP1 in presence of 30 μM ATP. Fluorescence imaging of 120 μM bpDNA mixing with (c) 10 μM eGFP-TNP1 or (d) 20 μM eGFP-TNP1.

Interestingly, by adding DNA molecules of 120 *μM/bp* into the 10*μM* eGFP-TNP1 solution (R_+/−_=0.5), phase-separated droplets appeared in the fluorescence images (Fig. 6c). As the content of TNP1 increases to 20 *μM* (R_+/−_ = 1.0), more and larger droplets showed up (Fig. 6d). This result indicates that the formation of TNP1-DNA droplets is not merely a consequence of charge neutralization. The polyelectrolyte properties, the bulky property (i.e., the size) of DNA as well as the special role of the phosphate group are all crucial influence factors on the formation of TNP1: DNA droplets.

## Discussion

The aim of our study is to examine the higher-order dynamics of TNP1 and DNA molecules. To achieve this, we employed a combination of computational and experimental techniques to characterize the interactions between TNP1 and DNA at both the atomistic and mesoscale levels. As a critical regulator of human genetic material preservation, TNP1 is characterized by its unique sequence composition of abundant positive charges and intrinsic disorder. The intrinsic disorder and multivalent properties pose challenges in the in vitro expression and purification of TNP1. To this end, we first attempted to purify nuclear transition proteins using ion exchange columns and a centrifuge method and were successful in obtaining the target proteins (Fig. S1).

We found that TNP1 binds strongly to DNA with a K_D_ of 0.9 uM (Fig. 1). The TNP1: DNA binding propensity is supposed to be stronger than histone: DNA binding, making it an effective substitute. However, it should not be too strong as it needs to be further replaced by protamine (Balhorn, 2007). In comparison to human protamine (24 arginines of 51 amino acids), TNP1 has a more balanced composition of 11 arginines and 10 lysines, suggesting that protamine could compete with TNP1 for binding to DNA. Further studies are needed to quantify this hypothesis. In addition, we observed that arginine can insert into the DNA grooves and interact with isolated O2’ atoms of thymine (Fig. 2). These kinds of interactions may have an influence on the DNA folding pattern and relate to the unwinding of DNA molecules or even local disruption of DNA chain (Rohs et al., 2009).

TNP1 binding reduces the stiffness of DNA, as revealed by AFM (Fig. 3), and helps DNA to adopt a more compact structure by reducing the energy cost of curling or looping. TNP1 also screens the electrostatic interactions between DNAs, contributing to the higher-order condensation of the TNP1-DNA complex. WCA potential-based computational simulation showed that the TNP1-DNA mixture quickly condensates into a single large cluster at high ratios of positive and negative charges (Fig. 4). Further studies with a larger number of TNP1 and DNA molecules are required to confirm this speculation. Due to the limitation of computational power, the computational simulation could only include limited number of TNP1 and DNA. But we could speculate that as more TNP1 and DNA are added to the system, the cluster would further grow up. AFM results showed that at higher concentrations of TNP1, the mixture of TNP1: DNA forms a denser hinged network with some of the DNA compressed into microspheres (Fig. 5), and fluorescence microscopy indicated the formation of phase-separated droplets in solution (Fig. 6). The size of the droplets depends on the concentration of TNP1.These results reflect the complicated dynamic behavior between TNP1 and DNA.

In summary, we investigated the molecular binding and condensation behavior between DNA and TNP1 at different concentrations. Our findings showed that TNP1 binding to DNA is influenced by electrostatic screening and elasticity changes, which aid the assembly of TNP1 with DNA. There are still many questions that need to be answered in the field of DNA condensation during spermatogenesis. For example, what are the concentration of TNP1 and the corresponding R_+/−_ at physiological conditions? Whether the TNP1 express at a low concentration that mainly aims to compete with histone or express at a high level that may condensate the DNA into a compact form? And how the TNP1 may be further replaced by protamine? These obscure questions are still yet to be explored. Additionally, salmon protamine has been used in recent years as a critical tool for mRNA delivery (Jarzebska et al., 2020; Jarzebska et al., 2021), however, its strong binding with nucleic acids makes it unfavorable for RNA release. In contrast, TNP1 is expected to have weaker binding with nucleotide acids than protamine, which makes it a potential candidate for DNA/RNA encapsulation. Therefore, our study not only provides fundamental knowledge for further understanding the role of TNP1 in histone-protamine replacement, but also gives insight into the field of vaccine development via nucleic acid – protein condensates.

## Methods

### Preparing DNA constructs

Experimentally, we used two kinds of DNA constructs, a short DNA of 177 bp and a long DNA of 3043 bp. The 177 bp DNA fragment contains the 601-positioning sequence. The long DNA contains a nucleotide sequence corresponding to the green fluorescence protein (GFP). We extracted the 3043 bp pUC75-eGFP Plasmid DNA, by using a commercial kit (TIANpure Mini Plasmid Kit, Beijing, China). The plasmid DNA was cut with BamH1 to a linear DNA (Takara Biomedical Technology (Beijing)). Finally, we measured the concentration and purity using a spectrophotometer (Thermo Fisher NanoDrop Lite; Waltham, MA). Samples with A260/A280 ratios of less than 1.7 were discarded.

### Cloning, Expression and purification of TNP1

The full-length Spermatid nuclear transition protein 1 (UniProtKB— P09430 · TNP1_HUMAN) sequence was cloned into a pET30a-MBP expression backbone (TEV protease cleavage site) and transformed into BL21 (DE3) competent cells. Inoculation of individual colonies were grown for overnight. Cultures were grown at 37°C until OD600 increasing to 0.5~0.6. The cultures were induced with 0.5 mM IPTG at 16°C for 16-18 hours for harvest. The cultures were collected by centrifugation and resuspended in a lysis buffer containing 1x PBS buffer pH 7.0, 1 mM PMSF, 2M Urea, 1mM DNAse, 10% Glycerol, 500 mM NaCl, and protease inhibitors (Calbiochem). Cell lysis was performed using sonication. The lysate was centrifuged at 18000 rpm for 20 min at 4°C.

The supernatant containing the MBP-TNP1 protein was then passed through Ni^2^+-NTA resin. The protein was eluted from the resin with the buffer containing 1x PBS buffer pH 7.0, 500 mM NaCl and 200 mM imidazole. Anion-exchange chromatography with a HiTrap S column (GE Healthcare Life Sciences) was further used to remove potent DNA/RNA contamination. The A260/280 ratio was measured below 0.5. The MBP-TNP1 proteins were cleaved overnight with homemade TEV protease to remove MBP tags and was isolated and concentrated by 10-kDa Amicon Ultra centrifugal filter unit. Filtered protein was confirmed with the SDS-PAGE, and its concentration was assessed by protein absorbance at 280 nm. Proteins were frozen in liquid N2 in aliquots and stored at −80 °C. The pET30a-MBP-EGFP-TNP1 expression and purification methods follows similar procedure as described above.

### Gel Electrophoresis

The samples of 177bp DNA (450 uM/bp) were mixed with TNP1 of different concentration (0–10 μM). The migration behavior of DNA on the 1% agarose gel under the action of protein was observed by electrophoresis.

### Isothermal titration calorimetry (ITC)

ITC experiments were carried out at 25 °C. TNP1 was titrated into the DNA solution of 2 uM/bp. The heat involved in the dilution of the TNP1 was measured by titration of TNP1 into PBS buffer. The dilution heat was subtracted from the total heat. The isothermal titration results were fitted to a one set of site model to give relevant thermodynamic parameters, including binding constant (K_D_), enthalpy change (ΔH). entropy change (-T ΔS), and the Gibbs free energy change (ΔG).

### Measurement of ζ-Potential

The zeta potential (ζ) of DNA-TNP1 complexes were determined using Zetasizer nano ZS90. The ζ-potential of the specimen was determined by recording the mobile velocity and direction of charged particles in an electric field. Analysis was performed in triplicate at 25 °C.

### CD spectrum

Circular dichroism experiments were carried using Chirascan Applied Photophysics spectropolarimeter. All the experiments were carried at room temperature with quartz cuvette of 1 mm path length. 177bp DNA at a fixed concentration of 450 μM was titrated with varied concentrations of TNP1 (0–10 μM) at pH 7, 1x PBS buffer. The measurements were taken at a wavelength range of 200 nm–300 nm and at a scan speed of 75 nm/min. 3 scans were recorded and averaged. Baseline spectra of a buffer was always subtracted from the spectra. The measurement was carried out by incubating the samples at 4 °C for 10 min.

### AFM sample preparation, data collection and analysis

Atomic force microscopy (Bruker) were used to image the DNA and DNA-TNP1 complex. Aliquots of 1–2 ml cut 3.043 kb pUC75-eGFP Plasmid DNA (1–10 ng/ml) in 5 mM MgCl_2_ were applied to freshly cleaved mica and allowed to bind for 2 min. The DNA solution concentration is about 1.5 uM/bp DNA. The mica was then rinsed five times with 1 ml distilled water and the excess liquids were wicked away at the mica edge with a tissue. To prepare samples of TNP1 and DNA mixture, we followed the procedure by Ukogu et al., 2020. A more dilute 0.3 uM/bp DNA concentration was used to prevent molecular clumping. The TNP1 concentration was controlled between 0.1– 1 uM TNP1. AFM Images were collected at room temperature with the constant force mode. The raw data images were analyzed using NanoScope Analysis software.

### Dynamic light scattering

The particle size of DNA-TNP1 complexes was measured with dynamic light scattering (DLS). For the preparation of DNA-TNP1 complexes with different R_+/_-ratios (0, 0.5, 1, 2), 100 *μ*M DNA was mixed with TNP1 at different concentrations and incubated for 5 mins for measurement. DLS analysis was performed in triplicate at 25 °C, scanning 10 times for each sample. The average % intensity and the dimeter of specimen was recorded for each sample.

### Fluorescence imaging of TNP1-DNA mixture

177bp DNA at a fixed concentration of 120 μM was mixed with EGFP-TNP1 of concentration between 0–20 μM at pH 7, 1x PBS buffer. 5 uL protein-DNA samples were plated on the glass slides slide and incubated (#1.5 glass thickness; Grace Bio-Labs) for a minimum of 10 minutes. Imaging was performed on a Zeiss Axio Observer 7 inverted microscope equipped with an LSM900 laser scanning confocal module and employing a 20x setting (Babinchak et al., 2020). eGFP was excited to fluoresce with a 488 nm laser.

### Conventional molecular dynamics simulation

We simulated the binding of TNP1 with DNA with all-atom dynamics simulations. The predicted structure of TNP1 was downloaded from UniProt and was firstly boiled at 600K to fully unfold the molecule. For the DNA molecule, we used a 40 bp short DNA segment with a sequence of 5’-AAGTACAAACTTTCTTGTATAAGTACAAACTTTCTTGTAT-3’, which was sourced from a synthetic DNA operator. The TNP1 was placed distal from the DNA molecule in a simulation box, so we could observe the spontaneous binding process between TNP1 and DNA. Each set of simulations was simulated for 100 ns and repeated 3 times. After the end of the simulations, we took out the final configuration of the TNP1-DNA complex and placed another TNP1 into the system, and in the same way simulate the binding of 2nd TNP1 to the formed TNP1-DNA complex. The cycle lasted 3 times till the 4th TNP1 was added to the system. All simulation systems were listed in Table S1. The simulations were performed with Gromacs 2019. We used standard simulation setup and parameters to perform the conventional dynamics simulation, briefly 300K for temperature, Lincs algorithm, for bond constraints, a cut-off of 1.0 nm for LJ interaction, and PME for electrostatic interaction. The MM-PBSA energy was estimated with the g_mmpbsa tool. A dielectric constant of 8 was used to estimate the PB energy and Coulomb interaction. The last 50 ns trajectories were used for MM-PBSA energy estimation(Wang et al., 2019).

### Sampling of TNP1-DNA configurations with metadynamcis

To enhance the sampling of TNP1-DNA binding configurations, we further performed metadynamics simulations. The collective variable was selected as the number of contact between phosphate (P) atom and nitrogen atoms of the arginine guanidinium group or the carbon atom of the lysine amine group. This would encourage the exploration of more contact between the phosphate group and guanidinium or amine group. A cut-off distance of 1.5 nm was used to judge the contact between P and N/C atoms. The DNA molecules were restrained during the metadynamics simulation. The widths and heights of the Gaussian hills were both set as 5. The metadynamics was performed for 20 ns. We performed 20 sets of simulation systems, starting from different initial displacement of TNP1 relative to DNA. After the metadynamics simulation, a 30ns conventional molecular dynamics simulation was followed to further relax and equilibrium the final configuration accessed via metadynamics.

### Assembly simulations of TNP1-DNA mixture

The Weeks-Chandler-Anderson (WCA) potential was used to simulate the assembly process of TNP1 and DNA (Mukherjee et al., 2021). DNA and TNP1 were both coarse-grained based on the paper of XX. Specifically, a DNA base pair was represented by 5 beads (PH1-SG1-BP-SG2-PH2). The centered BP has a diameter of sigma of 7.8 angstroms. SG1, SG2, PH1, and PH2 have a diameter of 4.2 angstroms. PH1 and PH2 each carry a negative charge. The neighboring-based pairs were connected and spatially constrained with bonds, angles, and dihedral potential. One hand, it maintains the rigidity of DNA at the principal direction. On the other hand, it maintains the surface helical contour (minor and major groove) of DNA molecules.

For protein TNP1, the protein is mapped to the CG model with the same approach as in the SCORPION mode (Ha-Duong, 2010). The main chain of TNP1 is consisted of BKB1 beads, while the side chain of amino acids typically consists of 1 to 3 beads. Arginine and lysine each consists of a positive charge. No rotation restraints were applied to the TNP1 peptide, giving full flexibility to TNP1 molecules. Detailed parameter set-up for DNA, TNP1, and counter ions (Na+ and Cl-) could refer to the study of Mukhejee and coworkers (Mukherjee et al, 2021a).

In a simulation box of 50 nm on each side, 30 DNA were mixed with 30, 60, 90, 120, and 180 DNA respectively in different simulations. A dielectric constant of 78.0 was applied to the system. Only counter ions of DNA and TNP1 are included the in the simulation system. PME was used for electrostatic interaction calculation. For the WCA potential, all interactions were cut-off at 2(sigma)^1/2^. The tabulated potential in Gromacs 2019 was applied to use the Weeks-Chandler-Anderson potential.

## Acknowledgements

This work was supported in party by Ministry of Science and Technology (2020YFA0908500 to S.Y.) and the National Natural Science Foundation of China (31971127 to S.Y). L.Z.L was supported by start-up funding.

## Author Contribution

**L.Z.L, X.S.Q, Y.S**: Conceptualization, **X.S.Q., Y.C.R.**: Data curation, Investigation, Writing- Original draft preparation. **L.Z.L**: Modeling, Visualization. ***Y.S.***: Supervision. **X.S.Q., L.Z. L., Y.S**: Writing- Reviewing and Editing.

