## Supplementary_Information for "Probing the hierarchical dynamics of DNA-sperm nuclear transition protein complex through fuzzy interaction and mesoscale condensation"

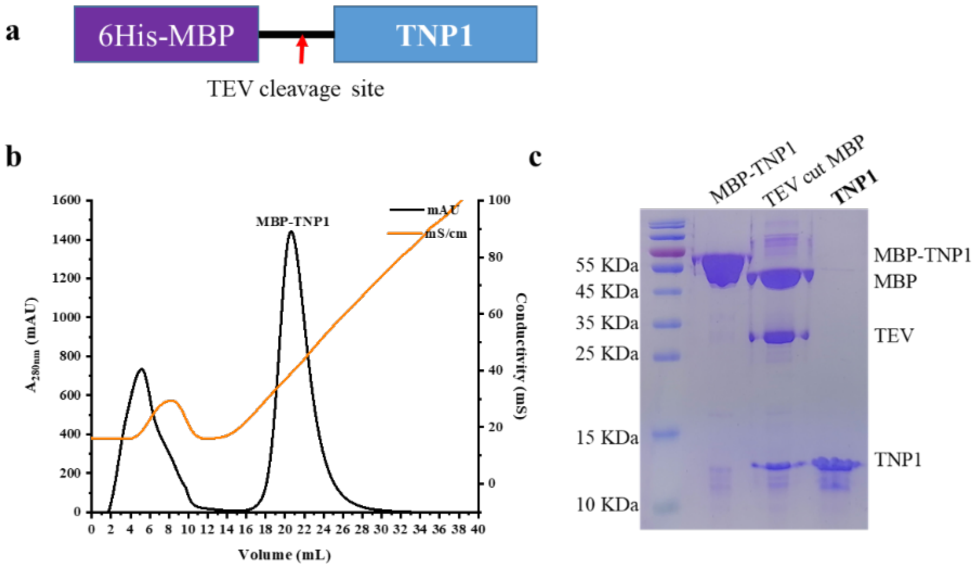


**Figure S1: Expression and Purification TNP1.** (a)Plasmid construct. (b) Anion-exchange chromatography with a HiTrap S column. (c) SDS-Page of MBP-TNP1,TEV cut MBP-tag and TNP1.

### Table S1: List of computational systems

| Conventional MD | | | | |
|---|---|---|---|---|
| | $1^{st}$ DNA Time | $2^{nd}$ DNA Time | $3^{rd}$ DNA Time | $4^{th}$ DNA Time |
| #1-#3 | 100 ns | 100 ns | 100 ns | 100 ns |

| MetaDynamics & Conventional MD | | |
|---|---|---|
| | Time | Time |
| #1-#20 | 20 ns | 30 ns |

| WCA potential based MesoMD | | |
|---|---|---|
| | Protein vs DNA | Time |
| #1 | 30: 30 | 100 ns |
| | 60: 30 | 100 ns |
| | 120: 30 | 100 ns |
| | 180: 30 | 100 ns |
| | 240: 30 | 100 ns |

#### First TNP1

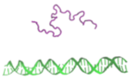

| | Van der Wall Energy | Electrostatic energy | Polar solvation Energy | SASA Energy | Binding Energy |
|---|---|---|---|---|---|
| #1 | -651.540 +/- 76.650 | -8829.372 +/- 181.557 | 4639.679 +/- 283.160 | -93.792 +/- 7.023 | -4935.025 +/- 79.020 |
| #2 | -378.791 +/- 52.874 | -8660.352 +/-198.607 | 3346.314 +/- 273.093 | -68.419 +/- 4.411 | -5461.248 +/- 179.771 |
| #3 | -324.959 +/- 39.065 | -8887.285 +/- 243.068 | 3932.756 +/- 347.091 | -62.877 +/- 5.524 | -5342.365 +/- 123.070 |
| Average | | | | | -5246.212 +/-275.97 |

#### Second TNP1

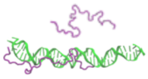

| | Van der Wall Energy | Electrostatic energy | Polar solvation Energy | SASA Energy | Binding Energy |
|---|---|---|---|---|---|
| #1 | -587.869 +/- 63.608 | -6999.452 +/-156.402 | 3992.172 +/- 297.726 | -73.851 +/- 6.181 | -3669.001 +/- 182.723 |
| #2 | -711.902 +/- 38.798 | -6996.487 +/-144.507 | 4092.904 +/- 279.643 | -86.490 +/- 4.466 | -3701.975 +/- 148.780 |
| #3 | -370.879 +/- 38.185 | -7013.783 +/- 117.377 | 3529.047 +/- 120.218 | -51.000 +/- 2.596 | -3906.615 +/- 77.901 |
| Average | | | | | -3759.197 +/-128.73 |

#### Third TNP1

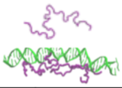

| | Van der Wall Energy | Electrostatic energy | Polar solvation Energy | SASA Energy | Binding Energy |
|---|---|---|---|---|---|
| #1 | -486.518 +/- 84.552 | -3891.004 +/- 546.778 | 1808.582 +/- 696.887 | -61.371 +/- 10.907 | -2630.310 +/- 269.427 |
| #2 | -462.690 +/- 56.904 | -4932.031 +/-117.716 | 3494.612 +/- 216.437 | -66.063 +/- 5.541 | -1966.173 +/- 116.028 |
| #3 | -628.160 +/- 83.321 | -5029.440 +/- 150.782 | 3765.859 +/- 334.494 | -84.525 +/- 7.614 | -1976.267 +/- 165.080 |
| Average | | | | | -2190.92 +/- 380.56 |

#### Fourth TNP1

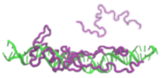

| | Van der Wall Energy | Electrostatic energy | Polar solvation Energy | SASA Energy | Binding Energy |
|---|---|---|---|---|---|
| #1 | -595.611 +/- 58.761 | -3160.143 +/-87.944 | 2844.414 +/- 231.137 | -74.129 +/- 7.279 | -984.469 +/- 168.636 |
| #2 | -362.525 +/- 52.674 | -2603.994 +/-113.947 | 2361.422 +/- 255.034 | -53.504 +/- 4.802 | -658.601 +/- 171.609 |
| #3 | -224.661 +/- 27.328 | -2287.626 +/- 47.519 | 1758.375 +/- 153.874 | -35.141 +/- 3.339 | -789.052 +/- 134.925 |
| Average | | | | | -810.71 +/- 164.01 |

**Figure S2**: MMPBSA energy for sequential binding of the first, second, third, and fourth TNP1 to a DNA molecule (40 bp). Solute dielectric constant is set as 8.0. Energy unit is KJ/mol.

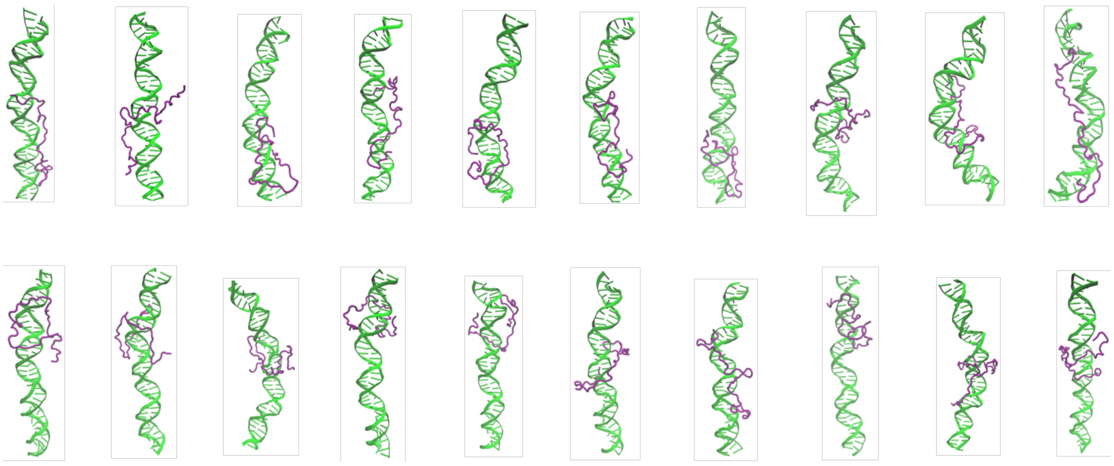

**Figure S3:** Final configurations of TNP1-DNA complex in 20 sets of MD simulations.

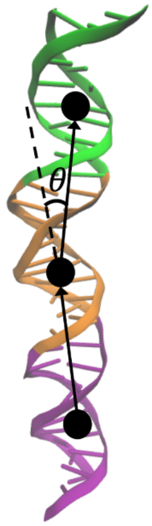


**Figure S4:** Definition of bending angle. A segment (marked in purple, orange, and green) consists of arbitrary 1o connected base pairs. The bending angle is defined as the cross angle between the vector connecting the (i-1)$^{th}$ and i$^{th}$ segment and the vector connecting the i$^{th}$ and (i+1)$^{th}$ segment.

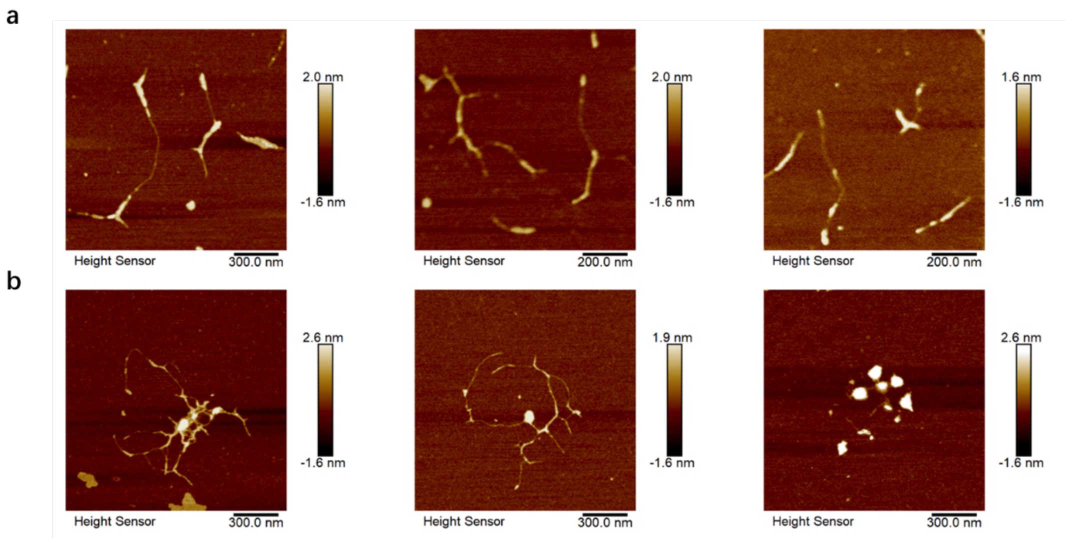


**Figure S5**: AFM images of (a) pure DNA (1.5 $\mu$M/bp) and (b) DNA-TNP1 complex (DNA of 0.3 $\mu$M/bp and TNP1 of 0.1 $\mu$M)

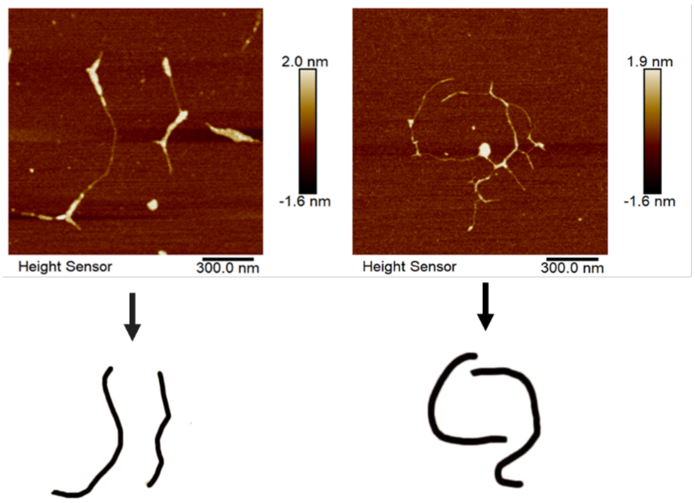


**Figure S6:** Extraction of DNA contour from AFM images.
